# Hyperactivity in mice induced by opioid agonists with partial intrinsic efficacy and biased agonism; alone and in combination with morphine

**DOI:** 10.1101/2023.05.11.540403

**Authors:** Agnes Acevedo-Canabal, Travis W. Grim, Cullen L. Schmid, Nina McFague, Edward L. Stahl, Nicole M. Kennedy, Thomas D. Bannister, Laura M. Bohn

**Author notes:** Correspondence: Laura M. Bohn.

## Abstract

Opioid analgesics like morphine and fentanyl induce mu-opioid receptor (MOR)-mediated hyperactivity in mice. Here we show that morphine, fentanyl, SR-17018, and oliceridine have submaximal intrinsic efficacy in the mouse striatum using ^35^S-GTPγS binding assays. While all of the agonists act as partial agonists for stimulating G protein coupling in striatum, morphine, fentanyl and oliceridine are fully efficacious in stimulating locomotor activity; meanwhile, the noncompetitive biased agonists, SR-17018 and SR-15099 produce submaximal hyperactivity. Moreover, the combination of SR-17018 and morphine attenuates hyperactivity while antinociceptive efficacy is increased. The combination of oliceridine with morphine increases hyperactivity which is maintained over time. These findings provide evidence that noncompetitive agonists at MOR can be used to suppress morphine-induced hyperactivity while enhancing antinociceptive efficacy; moreover, they demonstrate that intrinsic efficacy measured at the receptor level is not directly proportional to drug efficacy in the locomotor activity assay.

## Introduction

Opioid analgesics like morphine induce hyperlocomotion in addition to antinociception in rodents [1-3]; both effects result from MOR activation as demonstrated by a lack of response to opioid agonists in MOR-KO mice [4,5]. The MOR is a G protein-coupled receptor (GPCR) primarily known for activating inhibitory Gα_i/o_ proteins. It also interacts with scaffolding proteins such as βarrestins, to regulate and modulate receptor activity. Previous work using βarrestin2-knockout (βarrestin2-KO) mice showed that morphine induced greater antinociception in the absence of βarrestin2 with less development of tolerance [6,7]. The βarrestin2-KO mice also display less morphine-induced hyperactivity [8,9]. We described a series of structurally related MOR-selective agonists that display a preference for inducing MOR-G protein signaling over βarrestin2 recruitment (Schmid et al., 2017). From this series of compounds, SR-17018, was shown to produce antinociception with minimal respiratory suppression in mice [10,11] and Rhesus monkeys [12]. Moreover, chronic treatment with SR-17018 does not produce tolerance in the hot plate, formalin, and paclitaxel-induced neuropathic pain models in mice [13,14].

In this study, SR-15099 and SR-17018 were selected because they show a wide degree of “bias” between G protein signaling (^35^S-GTPγS binding) and βarrestin2 recruitment (enzyme fragment complementation, EFC) and because they are readily delivered to the brain following systemic injections and have a half-life over 6 hours in mice [11]. Recently, we showed these compounds are noncompetitive agonists at MOR using ^3^H-diprenorphine and ^3^H-naloxone radioligand binding assays and mouse brainstem GTPγS binding assays [15]. In that same study, we showed that oliceridine, which has been reported to display a preference for G protein signaling (cAMP accumulation studies) over βarrestin2 (EFC) recruitment [16], is competitive with DAMGO [15]. The present study was undertaken to determine whether G protein signaling-biased agonists produce hyperactivity and if co-administration with morphine would produce additive or competitive effects.

Herein, we find that the efficacy observed in G protein signaling (^35^S-GTPγS binding in striatum) does not predict the propensity to induce maximal hyperactivity. While they are as efficacious as morphine and fentanyl in striatal membrane GTPγS binding assays, SR-15099 and SR-17018 produce very little hyperactivity, in contrast to morphine and fentanyl. Moreover, despite having the lowest intrinsic efficacies of the group, oliceridine produces peak hyperactivity to a similar degree as fentanyl and morphine. In combination with morphine, low doses of oliceridine leads to increases in hyperactivity. However, SR-17018 attenuates morphine-induced hyperactivity while potentiating antinociceptive efficacy. Overall, we describe a paradoxical relationship wherein the noncompetitive G protein biased agonist, SR-17018, can differentially modulate morphine-induced behaviors *in vivo*.

## Materials and methods

### Animal Care and Use

Adult C57BL/6J and MOR-KO mice were purchased from The Jackson Laboratory (C57BL/6J Stock No: 000664 and MOR-KO ((B6.129S2-Oprm1^*tm1Kff*^/J) Stock No: 007559) and propagated by homozygous breeding. A total of 613 male C57BL/6J, 37 female C57BL/6J, and 44 male MOR-KO mice were utilized for the studies. All experiments used naïve adult mice of 10-16 weeks of age. For male mice, weights ranged between 23-35 grams and females ranged among 17-26 grams. Male and female C57BL/6J experiments were separated and presented accordingly where indicated. Approximately 80% of C57BL/6 mice were acquired from Jackson Labs while generating the remaining 20% in the lab vivarium space. Mice were group-housed (3-5 mice per cage) with 1/4” corncob bedding and maintained on a 12-hour light/dark cycle with food and water *ad libitum*. The number of mice used per treatment and experiment is included in the figure legends. The use of all mice was following the National Institutes of Health Guidelines for the Care and Use of Laboratory Animals with approval by The Scripps Research Institute Animal Care and Use Committee.

### Compounds

Morphine sulfate pentahydrate was acquired from the NIDA Drug Supply Program or purchased from Millipore Sigma or Spectrum Chemicals. Fentanyl citrate, naloxone hydrochloride dihydrate and dextroamphetamine (*d*-AMPH) hemisulfate were purchased from Millipore Sigma. Oliceridine hydrochloride (TRV-130) was purchased from MedKoo Biosciences. SR-17018 and SR-15099 as mesylate salts were synthesized at Scripps Research as previously described and validated by NMR for purity greater than 95%. DAMGO [D-Ala2, N-Me-Phe4, Gly5-ol]-Enkephalin (trifluoroacetate salt) was purchased from Tocris. For the *in vitro* studies, test compounds were prepared as 10 mM stocks in 100% dimethyl sulfoxide (DMSO, Thermo Fisher) and stored at - 20°C in 10 µL aliquots to avoid repeated freeze-thaw cycles, while DAMGO was prepared as 10 mM stocks in sterile water. SR-17018 is difficult to solubilize therefore, extra care was taken with DMSO to assure purity. Since DMSO is hygroscopic, it was aliquoted in glass bottles and kept at 4°C before use (pure DMSO is solid at 4°C).

### Drug solutions preparation

All experiments were conducted by investigators blinded to treatment assignments by a different experimenter. SR compounds were dissolved from powder immediately before use in vehicle 1:1:8 (10%DMSO (first):10%Tween-80 (second): then 80% sterile water) and administered intraperitoneally (I.P.). Conventional opioids and *d*-AMPH were either made in the 1:1:8 vehicle or saline, as indicated. Morphine sulfate, fentanyl citrate, oliceridine, naloxone, and *d*-AMPH solutions were prepared based on the salt weight of the drug, while the mesylate salts of the SR compounds dosing was adjusted for the free base weight as previously described [11,13]. All compounds were administered at a volume of 10 μL/g mouse weight. For co-administration studies, two consecutive injections on alternating sides were given.

### ^35^S-GTPγS binding assay in mouse striata

The striatal membrane G protein coupling assays were conducted as previously described [11,15,17] with minor modifications. Striata isolated from untreated C57BL/6J (10-16 weeks old) male mice were homogenized by tissue tearer and glass-on-glass homogenization in membrane buffer (10 mM Tris-HCl pH 7.4, 100 mM NaCl, 1 mM EDTA), passed through a 26-gauge needle (insulin syringe) eight times and resuspended in assay buffer (50 mM Tris-HCl pH 7.4,100 mM NaCl, 5 mM MgCl2,1 mM EDTA, and 20μM GDP). For each reaction, 2.5μg of membrane protein per well were incubated in assay buffer containing ∼0.1 nM of ^35^S-GTPγS and with increasing concentrations of agonist in a total volume of 200μL for 2 hours at room temperature. The reactions were terminated through filtration with GF/B filters (Perkin Elmer) using a 96-well plate harvester (Brandel). Filters were dried overnight, and radioactivity counts were measured using a TopCount NXT HTS microplate scintillation and luminescence counter (PerkinElmer). In all cases, assays were performed in duplicate or triplicate, and technical replicates were averaged into a single data point before combining assays between days to generate means with error. EC_50_, IC_50,_ and E_max_ values were determined via nonlinear regression in GraphPad Prism 9.0 software when convergence was obtained. Each n represents 1 mouse.

### Locomotor activity

The Versamax Animal Activity Monitoring System [20 × 20 cm^2^] (Accuscan Instruments, Columbus, OH) was used to assess open field locomotor activity as previously described [8]. The system consists of a photocell-equipped automated open field chamber contained inside sound mitigating boxes to record locomotor activity. The Versadat software (Accuscan Instruments, Columbus, OH) records the total number of beam breaks made per animal and these were collected in 5-minute intervals. Mice were individually placed into the activity monitoring boxes for 30 minutes to habituate (except where indicated) to the new environment and record basal activity. Animals were removed, injected, and immediately put back into the boxes to monitor opioid-induced locomotor activity for the times indicated.

### Opioid-opioid interactions in the acute thermal antinociceptive responses

Antinociceptive responses to thermal stimuli were determined according to previously published protocols [11]. Mice were placed in a Plexiglass chamber (16” tall, 8” in diameter) on a ceramic plate heated to 52 °C (hot plate test; Hot plate Analgesia Meter, Columbus Instruments). Basal nociceptive responses were determined by timing the amount of time until a mouse licks or flutters its fore- or hind-paws, rapidly steps, or jumps. Warm water-tail immersion (49 °C) assays were conducted immediately before the hot plate test. To avoid tissue damage, we imposed a ceiling time of 30 seconds for the tail-flick response and 20 seconds for the hot plate. Antinociceptive responses were collected at the indicated time points immediately following injection. Data are presented as “% maximum possible effect,” which was calculated by (response latency – baseline) / (maximal response cutoff latency – baseline) * 100%.

### Software and statistical analysis

Data are presented as the average of the mean ± SEM or as the mean with 95% conifidence intervals as indicated. Data were analyzed utilizing GraphPad Prism 9 (for Mac, GraphPad Software, San Diego, California USA, www.graphpad.com). For GTPγS binding, the radioactivity counts were normalized to baseline and maximum response produced by DAMGO and a 3-parameter nonlinear regression analysis was applied to obtain the efficacy and potency of the compounds. Two-way repeated measures ANOVA were used to compare time course dependent dose effects and ordinary one-way ANOVA followed by a Šídák’ s multiple comparison post-hoc analyses were used to compare total distance traveled for drug effect (sums). Two-way RM-ANOVA results are shown as data table inserts into figures, and the results of one-way ANOVA are indicated by symbols indicated in the figures. Significance was selected using alpha = 0.05.

## Results

### Evaluation of the intrinsic efficacy and competitive nature of partial agonists in mouse striatal membranes

DAMGO, an enkephalin analog, is a full agonist for stimulating ^35^S-GTPγS binding in mouse striatal membrane preparations; while morphine, SR-17018, and fentanyl produce submaximal stimulation (Figure 1A). Oliceridine is a known agonists at MOR, however, in this endogenous setting, stimulation could not be observed over basal background signaling. A low efficacy agonist will antagonize the effects of a full agonist down to the level of activation that the partial agonist produces; oliceridine (Figure 1B) and fentanyl (Figure 1C), act as classical competitive partial agonists. Although SR-17018 is also a partial agonist (48% DAMGO-stimulation) in striatum, it is noncompetitive, as it does not antagonize DAMGO’ s effects in striatal membranes (Figure 1D). We previously showed that none of these agonists stimulate GTPγS binding in MOR-KO mouse brainstem membranes [15]. Taken together with the prior study, SR-17018 acts as a noncompetitive partial agonist at MOR whereas oliceridine and fentanyl act as a competitive partial agonist in mouse striatum and brainstem.

**Figure 1.**
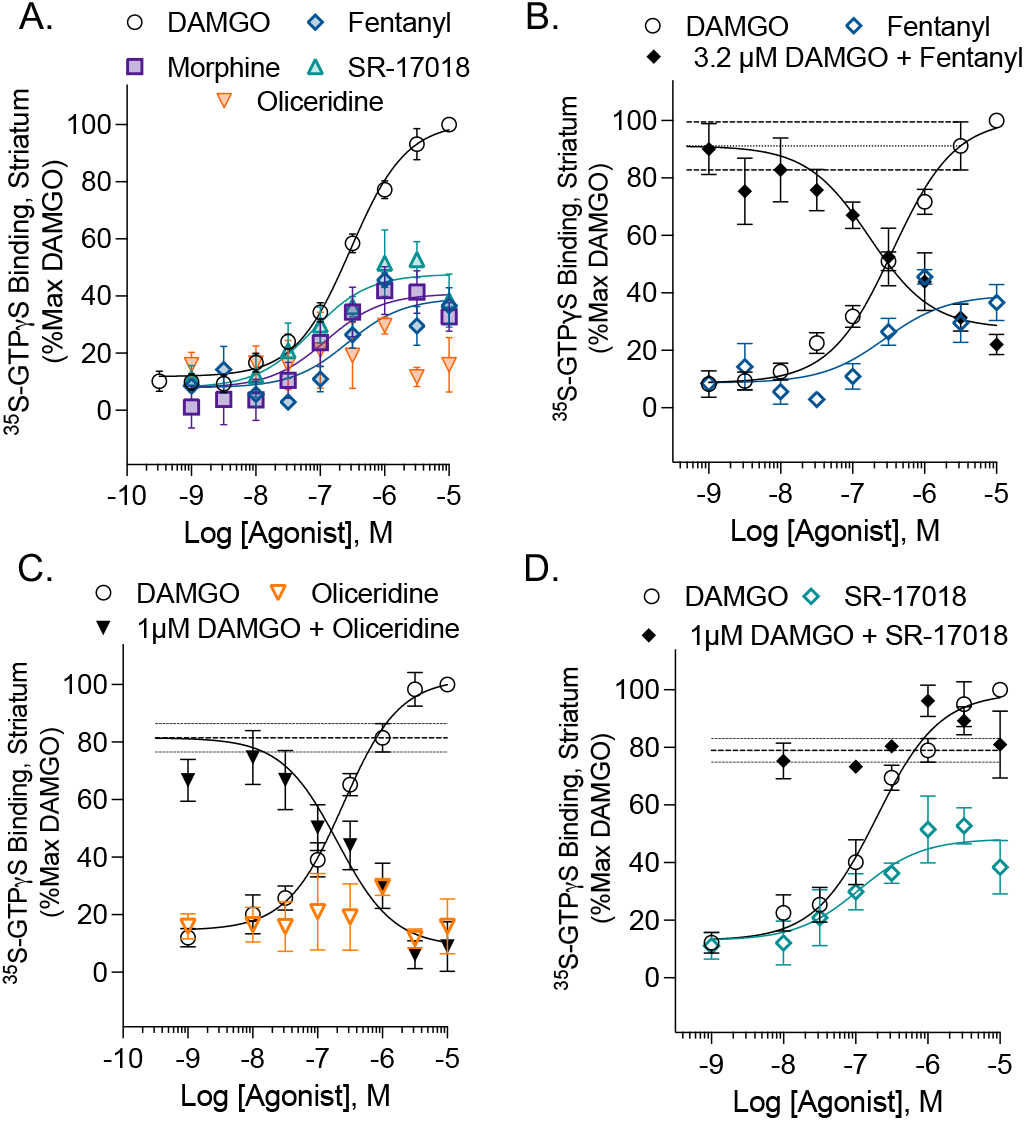
Evaluation of intrinsic efficacy of partial agonists at MOR in mouse striatum membranes by ^35^S-GTPγS binding. A. Morphine (Emax 41 (35-48)%; EC_50_:122 (50-298) nM, n = 9), fentanyl (Emax: 39 (31-47)%; EC_50_:151 (56-358) nM, n = 6); oliceridine (EC_50_: not converged, n = 4), and SR-17018 (Emax: 48 (48-58)%, EC_50_: 119 (105-128) nM, n = 5; are partial agonists relative to DAMGO (Emax: 100%, EC_50_ 290 (218-394) nM, n = 13). Partial agonist competition with full agonist activity can be observed for **B**. fentanyl (+ 3.2 µM DAMGO: IC_50_: 171 (50-520) nM, n = 3) and **C**. oliceridine (+ 1μM DAMGO: IC_50_ 203 (95% CI: 79-500) nM, n = 4) but not for **D**. SR-17018. DAMGO was run in parallel on each plate and IC_50_ calculations were constrained to the % stimulation obtained by DAMGO at the competitive dose on the plate; the mean of all the DAMGO curves is shown in A. The mean fold DAMGO stimulation was 1.41 ± 0.03 fold over baseline (1206 ± 252 cpm). Data are presented as mean ± SEM in the graphs and parameters as the mean with 95% CI; n = 1 refers to 1 mouse.

### Comparison of locomotor activity by different opioid agonists

Fentanyl, morphine, and oliceridine induce hyperactivity relative to vehicle (Figure 2A-C, interaction of time and treatment, p<0.0001) as previously reported [1-3,18]. SR-17018, and the related partial and biased noncompetitive agonist, SR-15099, also produce an increase in locomotor activity over time compared to vehicle (Figure 2D,E, interaction, p<0.0001). However, the modest stimulation induced by all doses of SR-15099 and SR-17018 is to a lesser extent than seen with the clinical analgesics which can be visualized by comparing the maximally efficacious dose (Figure 2F). Since the timing of peak drug effects can reflect differential pharmacokinetic properties (fentanyl and oliceridine having a faster onset and clearance relative to morphine, SR-17018, and SR-15099 [11,16], statistical analysis overtime was not applied. No hyperactivity is observed in MOR knockout (MOR-KO) mice at the highest doses tested (SFigure 1).

**Figure 2.**
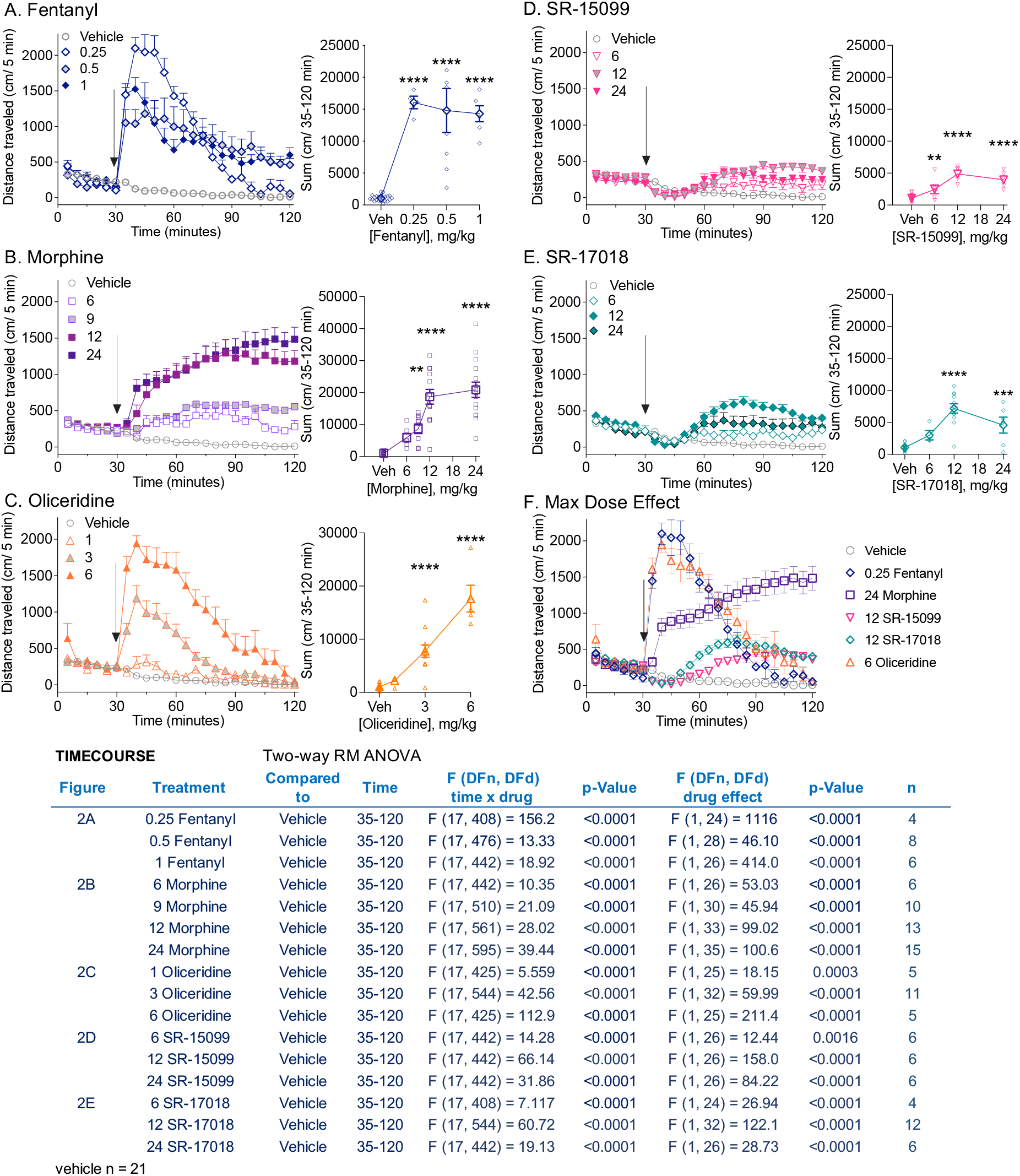
Comparison of locomotor activity by different opioid agonists. Locomotor activity using the open field locomotor boxes, measuring total distance traveled (cm/5-minute) with increasing doses of **A**. fentanyl, **B**. morphine, **C**. oliceridine, **D**. SR-15099, and **E**. SR-17018 in males C57BL6/J. A vehicle cohort is shown in each group for comparison. Basal locomotor activity was recorded for 30 minutes followed by injection with vehicle or test compound and activity was monitored for 90 additional minutes. All compounds were dissolved in vehicle 1:1:8 (10%DMSO:10%TWEEN80:80% sterile water). **A-E**. Time course of total distance traveled (cm/5-minute) (*left*) and total distance sums after treatment (cm/35-120 minutes) (*right*) are presented. Doses are indicated in the figure (mg/kg, i.p.). Two-way RM-ANVOA analysis for the time course data are presented in the statistical table inserted in the figure. The sum of distance traveled figures are shown as individual sums per mouse and mean ± SEM. For the sums, statistical comparison to vehicle was performed by ordinary one-way ANOVA followed by Dunnet’ s post-hoc comparison test (^**^p<0.01; ^***^p<0.001; ^****^p<0.0001). **F**. Time course comparison of the max dose effect of all tested compounds. Data are presented as mean ± SEM of the total distance (cm/5 min).

### Sensitization following chronic opioid exposure

Drug administration was continued in the mice from Figure 1 daily for 6 days at the indicated doses and mice were again assessed for opioid-induced locomotor activity on the seventh day at the same dose (Figure 3). As anticipated, fentanyl (Figure 3A) and morphine (Figure 3B) produce an enhanced response after chronic dosing (day effect: fentanyl: p=0.0004; morphine: p<0.0001). Since oliceridine (Figure 3C) has a more transitory effect, due to pharmacokinetic properties [18], analysis was compared for the first 30 min after injection; during that period, sensitization is also observed following chronic daily dosing (day effect: p=0.0461). Chronic treatment with SR-17018 (Figure 3D) and SR-15099 (Figure 3E) did not produce sensitization; moreover, SR-15099 daily administration led to a modest yet significant decrease in locomotor activity on day 7 (day effect: p=0.0179) over 90 minutes. Chronic vehicle treatment did not affect activity (Figure 3F). Individual mouse analysis for the sum of distance traveled is compared by one-way RM-ANOVA over 90 minutes (Figure 3G) and 30 minutes (Figure 3H) post-injection. Figure 3I shows that plasma and brain concentration of SR-15099 and SR-17018 do not differ between the first and seventh day of testing (unpaired two-tailed *t* test), suggesting that the compounds had access to mouse brain, although no sensitization occurred.

**Figure 3.**
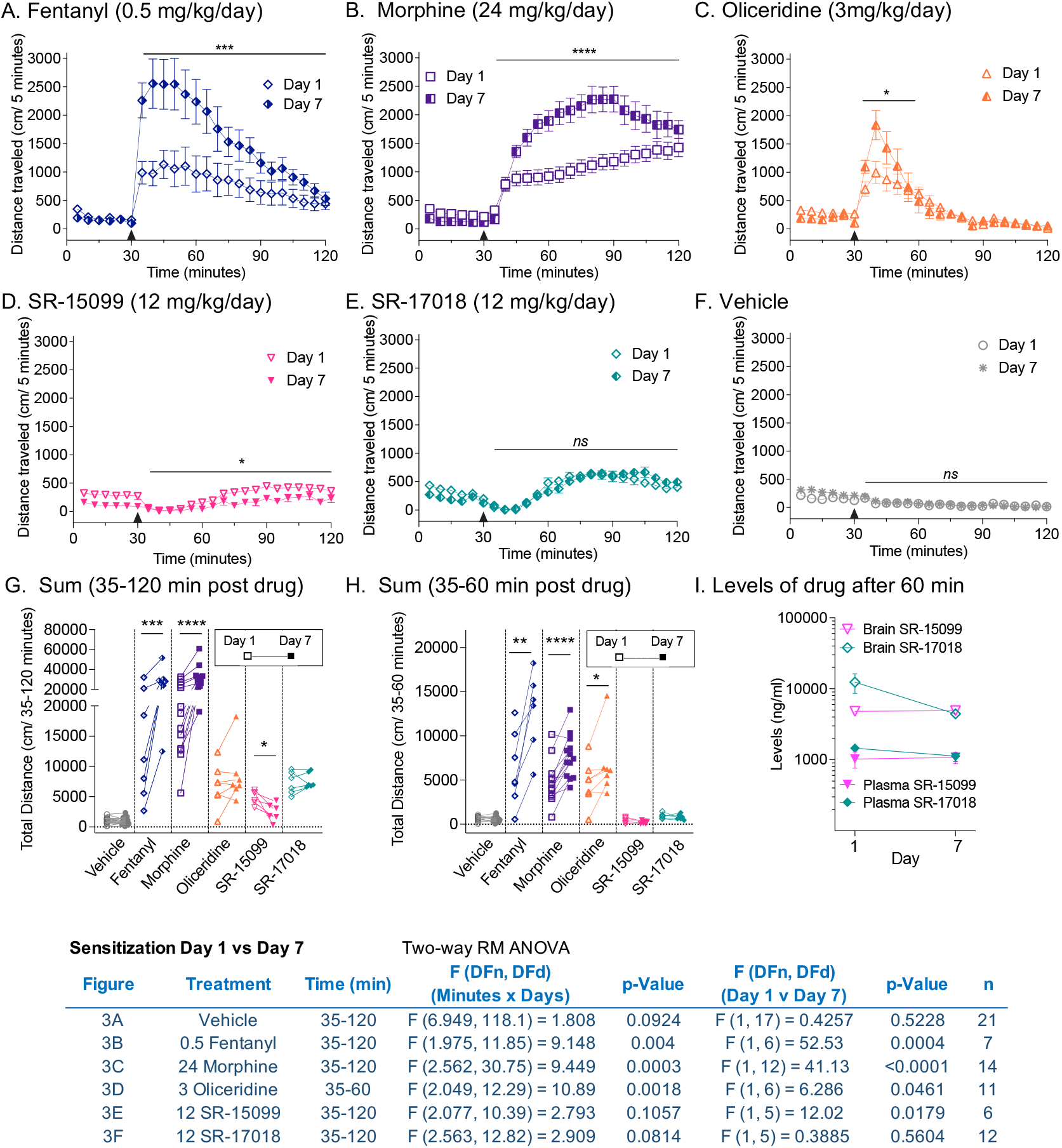
Repeated daily dosing of SR-17018 and SR-15099 does not lead to locomotor sensitization observed with fentanyl, morphine and oliceridine. Mice from Figure 2 were treated with daily doses of **A**. Fentanyl (0.5 mg/kg, i.p.); **B**. Morphine (24 mg/kg, i.p.); **C**. Oliceridine (3 mg/kg, i.p.); **D**. SR-15099 (12 mg/kg, i.p.); **E**. SR-17018 (12 mg/kg, i.p.) or **F**. Vehicle. On day 7, mice were i.p. challenged with the same dose and locomotor activity was assessed over 90 min following a 30 min habituation. The data are presented as mean ± SEM of the total distance (cm/5 min); **C**, Cumulative distance traveled sums (over 35-120 minutes) or **D**, (over 35-60 minutes) after treatment comparing Day 1 vs Day 7. Statistical comparison over time using 2-way RM-ANOVA is indicated in **A-F** (see inserted table for statistics); and for **G-H** by paired Student’ s t-test (^*^p<0.05; ^**^p<0.01 ^***^p<0.001; ^****^p<0.0001). **I**. Brain and plasma levels of SR-17018 and SR-15099 on day 1 and day 7 of daily dosing (12 mg/kg, i.p.) taken at 60 minutes after injection.

### SR-17018 attenuates morphine-induced locomotor activity

Since the peak effect of SR-17018 plateaus at a submaximal level relative to morphine, we asked what impact the combination of SR-17018 and morphine would have on mouse ambulatory behaviors. Given that the SR-17018 brain distribution peaks between 30-120 minutes [11,13], mice were pretreated 30 minutes prior to morphine administration with SR-17018 or vehicle. SR-17018 (6 mg/kg, I.P.) had no discernable impact on morphine (6 mg/kg, i.p.)-induced hyperactivity compared to morphine with vehicle pretreatment (Figure 4A, drug effect, p=0.2550, Figure 4B for comparison of total distance). At higher doses of morphine, SR-17018 dose-dependently attenuates locomotor activity induced by morphine (Figure 4C-H and see Table insert for a summary of pretreatment effect on second treatment statistical analyses). Notably, the combination of SR-17018 and morphine produces a sustained lower level of hyperactivity that remains consistently higher than activity produced in vehicle + saline-treated controls (Figures 4B, E H: SR-17018 + morphine vs vehicle + saline, ordinary one-way ANOVA: p<0.0001).

**Figure 4.**
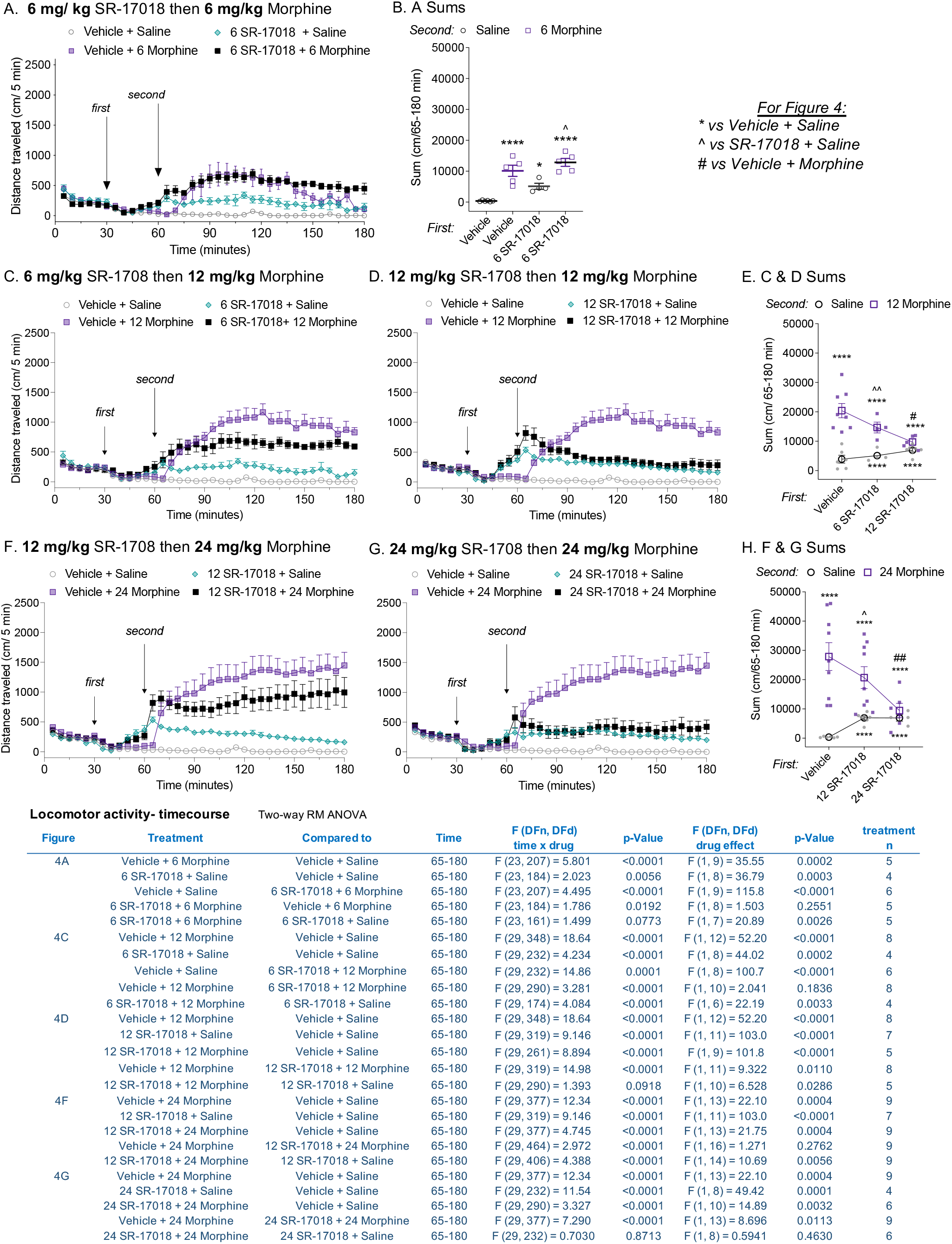
Pretreatment with SR-17018 dose-dependently attenuates morphine-induced locomotor activity. Mice were habituated for 30 minutes then given a 30 minute pretreatment *(first)* with vehicle or SR-17018 prior to saline or morphine challenge *(second)*; doses of each drug (IP) are indicated in the subheadings for each figure. **A**. SR-17018 (6 mg/kg) given before 6 mg/kg morphine time course and **B**. Sum of total distance travelled presented as individual mice and mean ± SEM. SR-17018 given in the first injection at **C**. 6 mg/kg or **D**. 12 mg/kg before 12 mg/kg morphine; **E**, sums for C and D. SR17018 given in the first injection at **F**. 12 mg/kg and **G**. 24 mg/kg before 24 mg/kg morphine; **H**. shows the sums for F & G. Data are presented as raw locomotor activity date with mean ± SEM of the total distance (cm) for A,C,D,E and F; and as the individual mouse sums for B, E and H with mean ± SEM. The inserted table presents the comparison of treatment effects over time by two-way RM-ANOVA; the sums are compared by ordinary one-way ANOVA (^*^p<0.05; ^****^p<0.0001 vs. vehicle + saline; ^#^p<0.05, ^##^p<0.01, ^####^p<0.0001 vs. vehicle + morphine and ^p<0.05, ^p<0.01 vs. SR-17018 + saline).

To further investigate the nature of the inhibitory effect, we tested whether SR-17018 could modulate activity produced by a psychostimulant, *d*-amphetamine. A combination of opioids and psychostimulants typically potentiates hyperactivity in mice [19,20] due to the combination of converging mechanisms that induce dopamine release [20,21]. As anticipated, morphine pretreatment enhances *d*-amphetamine-induced locomotor activity compared to vehicle pretreatment (SFigure 2A, interaction of time and drug: p<0.0001); pretreatment with SR-17018 also enhances amphetamine-induced hyperactivity (SFigure 2B, interaction of time and drug; p<0.0001). These observations suggest that SR-17018 does not act by a MOR-independent means of sedating mice and that it can still potentiate the effects of a psychostimulant, like morphine.

When administered 30 or 60 minutes after morphine, SR-17018 decreases morphine-induced hyperactivity (SFigure 3A-B, two-way RM-ANOVA interaction of time and dose at 30 mins: p=0.0002; and at 60 mins: p=0.0032 post treatment relative to morphine + vehicle; see table in SFigure 3 for post-hoc analysis). In all cases, SR-17018 decreases morphine-induced hyperlocomotion to a level that approaches stimulation by SR-17018 alone (SFigure 3A-B). On the other hand, naloxone fully antagonizes morphine-induced locomotor activity to resemble saline + saline-induced activity levels (SFigure 3C, two-way RM-ANOVA, interaction of drug and time: morphine + saline vs morphine + naloxone: p<0.0001; morphine + naloxone vs saline + saline, p=0.1106). This demonstrates an important difference between SR-17018 and naloxone, where naloxone serves as a competitive antagonist and blocks all of morphine’ s effects, SR-17018 appears to act as a partial agonist in vivo, where its own activity is maintained while tempering the effects of morphine.

### Co-treatment of partial agonists differentially effects morphine-induced hyperactivity and antinociception

Oliceridine and SR-17018 act as partial agonists in mouse striatum (Figure 1); however, only oliceridine acts as an orthosteric partial agonists as it competes with DAMGO-stimulated ^35^S-GTPγS binding in the striatal membranes. Furthermore, we had previously shown this same pharmacological profile in mouse brainstem (a region involved in descending pain perception) [15]. Therefore, we asked how the two agonists would interact with morphine in both the locomotor activity assay and the hot plate antinociception assay. Doses of each drug used were chosen based on producing a submaximal response in both the locomotor activity assay and the hot plate nociception assay, to allow for the observation of an additive effect. Oliceridine has a rapid onset and relatively short metabolic life in mice [16,18]; therefore, we opted for a co-treatment approach.

As expected from the pre- and post- treatment studies (Figures 4 and SFigure 3), co-treatment with SR-17018 suppresses morphine-induced hyperactivity (Figure 5A) (drug effect: vehicle + morphine vs SR-17018 + morphine, p=0.0123) to nearly the level of activity induced by SR-17018 alone. If the comparison is made following 60 minutes after injection, allowing for SR-17018 drug delivery the brain, the treatment groups do not differ (p= 0.2923), in agreement with the pretreatment studies (Figure 4). The combination of equi-efficacious doses of morphine and SR-17018 leads to an increase in antinociceptive response compared to the effect of each drug alone in the hot plate assay (Figure 5B, drug effect: SR-17018 + morphine: vs. morphine + vehicle, p=0.0019; vs saline + SR-17018, p=0.0001). This same effect was observed in female C57BL6 mice, wherein SR-17018 attenuates morphine-induced locomotor activity while enhancing morphine-induced antinociception (SFigure 5).

**Figure 5.**
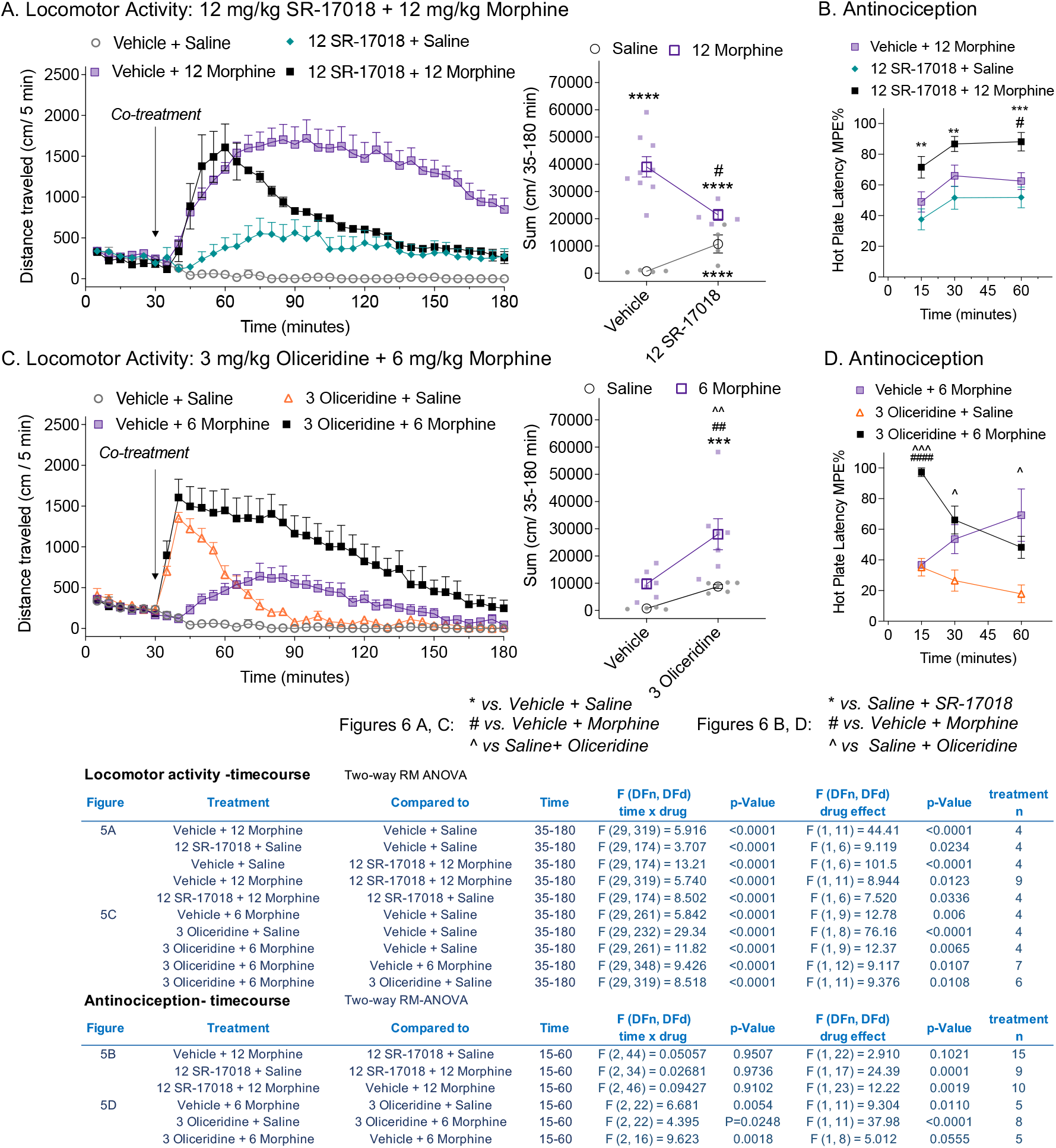
Co-treatment of SR-17018 with morphine decreases morphine-induced hyperactivity without attenuating antinociception. Comparing *in vivo* drug interactions in the open field locomotor activity assay showing total distance traveled (cm/5-minutes) (*left)* with total distance sums after co-treatment (cm/35-180 minutes) (*right*) and antinociception assessing the latency to respond to the hot plate (52°C) assay. **A**. Distance traveled over time following vehicle or SR-17018 (12 mg/kg, i.p.) co-administered with saline or morphine (12 mg/kg, i.p.) as indicated in the figure legend. **B**. Hot plate response latency following the same drug and vehicle combinations tested over 1 hour at the times indicated (mean ± SEM). Vehicle + saline was not tested in hot plate assay. **C**. Distance traveled over time following vehicle or oliceridine (3 mg/kg, i.p.) co-administered with saline or morphine (6 mg/kg, i.p.). **D**. Hot plate response latency. For the distance traveled (A, C), individual mice and the mean ± SEM are shown; statistical comparison by two-way RM-ANOVA is presented in the inserted table. The sums of the distance traveled are compared by ordinary one-way ANOVA (^#^p<0.05, ^##^p<0.01 vs. vehicle + morphine; ^***^p<0.001; ^****^p<0.0001 vs. vehicle + saline; ^p<0.01 vs. SR-17018 + saline). For the antinociception studies (B, D) analysis over time is presented in the antinociception -time course table with two-way RM-ANOVA; a Šídák’ s multiple comparisons post-hoc analysis determines significance: for C: ^**^p<0.01, ^***^p<0.001 vs. vehicle + morphine and ^#^p<0.05 vs. SR-17018 + saline and for D: ^p<0.05, ^^^^^p<0.001 vs. saline + oliceridine and ^####^p<0.0001 vs. vehicle + morphine.

Oliceridine administered with morphine produces more hyperactivity than with either drug alone (Figure 5C, drug effect: oliceridine + morphine: vs. vehicle + morphine, p=0.0.0107; vs oliceridine + saline, p=0.0108). Oliceridine significantly increases morphine-induced antinociception in the hot plate assay at the 15-minute timepoint (Figure 5D); however, over time, the effect of oliceridine appears to wear off, in agreement with its short-lived bioactivity. Notably, however, the enhanced locomotor activity persists after one hour past injection, although oliceridine’ s effects when administered with saline have waned by then (Figure 5C drug effect from 90-180 min comparing oliceridine + morphine: vs. vehicle + morphine, p=0.0497; vs. oliceridine + saline, p=0.0108).

## Discussion

In this study, we explore partial agonism *in vivo* and *in vitro* using MOR-dependent behaviors and mouse striatal membranes, respectively. In mouse striatal membranes, all the compounds tested, including morphine and fentanyl, act as partial agonists for inducing ^35^S-GTPγS binding (Figure 1), although their ability to promote hyperactivity varies. Oliceridine is a potent partial agonist *in vitro* that competes with DAMGO-stimulated ^35^S-GTPγS binding in brain (current study and [15] for brainstem) yet it also produces hyperactivity to the same extent as morphine and fentanyl. When oliceridine and morphine are combined at submaximal doses, locomotor activity and antinociception are both enhanced. SR-17018 is also a partial agonist in brain; however, it is noncompetitive with DAMGO-stimulated ^35^S-GTPγS coupling (current study and [15] for brainstem). In the locomotor activity assay, SR-17018 produces a blunted hyperactivity response and repeated dosing does not produce sensitization. The combination of SR-17018 and morphine decreases the hyperactivity produced by morphine while enhancing the antinociception observed.

Fentanyl, oliceridine, SR-17018, and SR-15099 have previously been described as biased agonists; where fentanyl has been shown to preferentially promote MOR-mediated βarrestin2 recruitment over G protein signaling while the other compounds are G protein signaling biased [11,16]. In the mouse brain, all of the compounds tested, including morphine and fentanyl, act as partial agonists for inducing G protein signaling (^35^S-GTPγS binding) (Figure 1 and [11,15,22-24]). Recently, it has been suggested that the improved therapeutic window (antinociception with less respiratory suppression) produced by oliceridine and SR-17018 is due to their low intrinsic efficacy at MOR as opposed to G protein signaling bias (Gillis et al., 2020). However, while fentanyl and morphine may appear to be highly efficacious agonists in frequently used receptor overexpression cellular systems; they are partial agonists with ∼50% efficacy when tested in mouse brain tissue where there is low receptor reserve (Figure 1 for striatum, [11] and [15] for brainstem). Therefore, low intrinsic efficacy alone does not account for the differences in physiological responses produced by these opioid agonists.

In biochemical measures of intrinsic activity, partial agonists will compete with a full agonist for receptor occupancy; as a function of dose, the submaximal stimulation induced by the partial agonist will overcome the response produced by the full agonist as the partial agonist reaches full occupancy of the receptor population. As increasing concentrations of a partial agonist compete for occupancy, the signal produced by the full agonist will decrease and the partial agonist will appear to have antagonistic properties. In behavioral studies, this is rarely observed with opioids, as combination of two partial agonists such as fentanyl and morphine, will generally lead to an additive or synergistic effect [25]. One exception may be buprenorphine, which has antagonistic properties in the presence of other opioid agonists such as fentanyl and morphine) depending on the response and the system. For example, buprenorphine produces a bell-shaped antinociceptive response in mice [26,27].; however, buprenorphine also has multiple receptor targets and it becomes difficult to fully explain these effects by actions at the MOR alone [28].

The comparison of locomotor activity profiles reveals a dramatic difference between the SR-17018 and SR-15099 biased agonists compared to the other compounds. While they are not devoid of producing hyperactivity, the profile captures an initial dip in activity with a restoration of activity by 60 minutes post injection; this ambulatory behavior is remarkably steady over time, and while it is greater than observed following vehicle treatment, it does not nearly reach the heights produced by the other agonists (Figure 2). In contrast, despite also being a G protein signaling biased agonist, oliceridine produces a different profile, with rapid-onset hyperactivity reaching peak effects similar to fentanyl (current study and [18]). Further, repeated dosing of fentanyl, morphine and oliceridine produces sensitization (Figure 3), while SR-17018- and SR-15099-induced activity remains consistent with chronic administration (with a modest decrease in activity observed for SR-15099). Notably, prior analysis of SR-17018 and SR-15099 in hot plate and tail flick nociceptive tests demonstrated that they reach the same efficacy as fentanyl and morphine, yet they do not produce respiratory suppression at doses exceeding their antinociceptive efficacy (48 mg/kg, i.p.) [11].

Locomotor hyperactivity and respiratory suppression are both affected by dopamine levels, which are elevated in response to typical opioid agonists [21,29]. It will be of interest to compare regional dopamine levels following the administration of these agonists in future studies. At high doses, opioids lead to skeletal muscle rigidity in mice and humans and this effect may contribute to overdose cases of respiratory failure [30,31]. Further, we recognize that high doses of opioids can lead to stiffness which could impact our ability to measure the animal’ s total distance traveled; a plateau effect can be observed for fentanyl in Figure 1, where a peak effect is observed at 0.25 mg/kg with higher doses producing no gain in distance.

Future studies are needed to delineate the rewarding properties of the novel SR G protein biased MOR ligands; however, recent studies have shown that SR compounds are not devoid of conditioned place preference potential [32]. Their poor solubility limits their utility in self-administration studies, although low doses were attempted by Dr. Marc Caron in the mouse self-administration paradigm. Given the low dose, the lack of observed self-administration was not very telling regarding the abuse potential of the compound (unpublished observations). The results from this current study may provide further insight on the pharmacological interactions of novel MOR compounds with morphine. Upon chronic administration, SR-17018 produces no tolerance in the hot plate, formalin and paclitaxel-induced neuropathy pain tests [13,14] and no sensitization to the locomotor stimulatory effects (Figure 3). Further, treatment of morphine-tolerant mice with SR-17018 suppresses withdrawal symptoms and restores morphine sensitivity in the hot plate antinociception [13,32]; the ability to blunt morphine-induced psychomotor activity may also prove to be beneficial as an abuse deterrent. Further studies are needed to explore the potential of these novel compounds.

## Supplementary Materials

SFigure 1: A lack of opioid-induced hyperactivity in MOR-KO mice; SFigure 2: SR-17018, like, morphine, increases d-amphetamine-induced locomotor activity. SFigure 3: SR-17018 reversal of morphine-induced locomotor activity; SFigure 4: Co-treatment of morphine with SR-15099 attenuates morphine-induced hyperactivity; SFigure 5: In female mice, co-treatment of morphine with SR-17018 decreases morphine-induced hyperactivity without attenuating antinociception.

## Author Contributions

Contributed to experimental design, performed the majority of experiments and analysis and wrote and revised the manuscript: AAC; Contributed to the experiment design and execution: TWG, CLS and ELS; experimental assistance: NM; synthesized SR-17018 and SR-15099: NMK; supervised chemical synthesis: TDB; wrote and revised the manuscript and performed analysis: LMB.

## Institutional Review Board Statement

Studies were conducted under the approved animal protocol 15-004 for LMB through the University of Florida Scripps Research IACUC.

## Data Availability Statement

Data are presented within the article and supplementary material. Raw values are plotted in the figures and are available upon request.

## References

1. Rethy, C.R.; Smith, C.B.; Villareal, J.E. Effects of Narcotic Analgesics Upon the Locomotor Activity and Brain Cathecolamine Content of the Mouse. Journal of Pharmacology and Experimental Therapeutics 1970, 176, 472–479.

2. Bailey, A.; Metaxas, A.; Al-Hasani, R.; Keyworth, H.L.; Forster, D.M.; Kitchen, I. Mouse strain differences in locomotor, sensitisation and rewarding effect of heroin; Association with alterations in MOP-r activation and dopamine transporter binding. European Journal of Neuroscience 2010, 31, 742–753, doi:10.1111/j.1460-9568.2010.07104.x.

3. Varshneya, N.B.; Walentiny, D.M.; Moisa, L.T.; Walker, T.D.; Akinfiresoye, L.R.; Beardsley, P.M. Opioid-like antinociceptive and locomotor effects of emerging fentanyl-related substances. Neuropharmacology 2019, 151, 171–179, doi:10.1016/j.neuropharm.2019.03.023.

4. Hall, F.S.; Li, X.F.; Goeb, M.; Roff, S.; Hoggatt, H.; Sora, I.; Uhl, G.R. Congenic C57BL/6 mu opiate receptor (MOR) knockout mice: baseline and opiate effects. Genes Brain Behav 2003, 2, 114–121, doi:10.1034/j.1601-183x.2003.00016.x.

5. Sora, I.; Elmer, G.; Funada, M.; Pieper, J.; Li, X.F.; Hall, F.S.; Uhl, G.R. Mu opiate receptor gene dose effects on different morphine actions: evidence for differential in vivo mu receptor reserve. Neuropsychopharmacology 2001, 25, 41–54, doi:10.1016/S0893-133X(00)00252-9.

6. Bohn, L.M.; Lefkowitz, R.J.; Gainetdinov, R.R.; Peppel, K.; Caron, M.G.; Lin, F.-t. Enhanced Morphine Analgesia in Mice Lacking Beta-Arrestin 2. 1999, 286, 2495–2499.

7. Bohn, L.M.; Gainetdinov, R.R.; Lin, F.T.; Lefkowitz, R.J.; Caron, M.G. Mu-opioid receptor desensitization by beta-arrestin-2 determines morphine tolerance but not dependence. Nature 2000, 408, 720–723, doi:10.1038/35047086.

8. Bohn, L.M.; Gainetdinov, R.R.; Sotnikova, T.D.; Medvedev, I.O.; Lefkowitz, R.J.; Dykstra, L.A.; Caron, M.G. Enhanced rewarding properties of morphine, but not cocaine, in beta(arrestin)-2 knock-out mice. J Neurosci 2003, 23, 10265–10273.

9. Mittal, N.; Tan, M.; Egbuta, O.; Desai, N.; Crawford, C.; Xie, C.W.; Evans, C.; Walwyn, W. Evidence that behavioral phenotypes of morphine in beta-arr2 -/- mice are due to the unmasking of JNK signaling. Neuropsychopharmacology 2012, 37, 1953–1962, doi:10.1038/npp.2012.42.

10. Gillis, A.; Gondin, A.B.; Kliewer, A.; Sanchez, J.; Lim, H.D.; Alamein, C.; Manandhar, P.; Santiago, M.; Fritzwanker, S.; Schmiedel, F.; et al. Low intrinsic efficacy for G protein activation can explain the improved side effect profiles of new opioid agonists. Sci Signal 2020, 13, doi:10.1126/scisignal.aaz3140.

11. Schmid, C.L.; Kennedy, N.M.; Ross, N.C.; Lovell, K.M.; Yue, Z.; Morgenweck, J.; Cameron, M.D.; Bannister, T.D.; Bohn, L.M. Bias Factor and Therapeutic Window Correlate to Predict Safer Opioid Analgesics. Cell 2017, 171, 1165–1175 e1113, doi:10.1016/j.cell.2017.10.035.

12. Cornelissen, J.C.; Blough, B.E.; Bohn, L.M.; Negus, S.S.; Banks, M.L. Some effects of putative G-protein biased mu-opioid receptor agonists in male rhesus monkeys. Behav Pharmacol 2021, doi:10.1097/FBP.0000000000000634.

13. Grim, T.W.; Schmid, C.L.; Stahl, E.L.; Pantouli, F.; Ho, J.H.; Acevedo-Canabal, A.; Kennedy, N.M.; Cameron, M.D.; Bannister, T.D.; Bohn, L.M. A G protein signaling-biased agonist at the mu-opioid receptor reverses morphine tolerance while preventing morphine withdrawal. Neuropsychopharmacology 2019, doi:10.1038/s41386-019-0491-8.

14. Pantouli, F.; Grim, T.W.; Schmid, C.L.; Acevedo-Canabal, A.; Kennedy, N.M.; Cameron, M.D.; Bannister, T.D.; Bohn, L.M. Comparison of morphine, oxycodone and the biased MOR agonist SR-17018 for tolerance and efficacy in mouse models of pain. Neuropharmacology 2021, 185, 108439, doi:10.1016/j.neuropharm.2020.108439.

15. Stahl, E.L.; Schmid, C.L.; Acevedo-Canabal, A.; Read, C.; Grim, T.W.; Kennedy, N.M.; Bannister, T.D.; Bohn, L.M. G protein signaling-biased mu opioid receptor agonists that produce sustained G protein activation are noncompetitive agonists. Proc Natl Acad Sci U S A 2021, 118, doi:10.1073/pnas.2102178118.

16. DeWire, S.M.; Yamashita, D.S.; Rominger, D.H.; Liu, G.; Cowan, C.L.; Graczyk, T.M.; Chen, X.T.; Pitis, P.M.; Gotchev, D.; Yuan, C.; et al. A G protein-biased ligand at the muopioid receptor is potently analgesic with reduced gastrointestinal and respiratory dysfunction compared with morphine. J Pharmacol Exp Ther 2013, 344, 708–717, doi:10.1124/jpet.112.201616.

17. Zhou, L.; Stahl, E.L.; Lovell, K.M.; Frankowski, K.J.; Prisinzano, T.E.; Aube, J.; Bohn, L.M. Characterization of kappa opioid receptor mediated, dynorphin-stimulated [35S]GTPgammaS binding in mouse striatum for the evaluation of selective KOR ligands in an endogenous setting. Neuropharmacology 2015, 99, 131–141, doi:10.1016/j.neuropharm.2015.07.001.

18. Mori, T.; Takemura, Y.; Arima, T.; Iwase, Y.; Narita, M.; Miyano, K.; Hamada, Y.; Suda, Y.; Matsuzawa, A.; Sugita, K.; et al. Further investigation of the rapid-onset and short-duration action of the G protein-biased mu-ligand oliceridine. Biochem Biophys Res Commun 2021, 534, 988–994, doi:10.1016/j.bbrc.2020.10.053.

19. Mori, T.; Ito, S.; Narita, M.; Suzuki, T.; Sawaguchi, T. Combined effects of psychostimulants and morphine on locomotor activity in mice. J Pharmacol Sci 2004, 96, 450–458, doi:10.1254/jphs.fpj04039x.

20. Trujillo, K.A.; Smith, M.L.; Guaderrama, M.M. Powerful behavioral interactions between methamphetamine and morphine. Pharmacol Biochem Behav 2011, 99, 451–458, doi:10.1016/j.pbb.2011.04.014.

21. Di Chiara, G.; Imperato, A. Drugs abused by humans preferentially increase synaptic dopamine concentrations in the mesolimbic system of freely moving rats. Proceedings of the National Academy of Sciences of the United States of America 1988, 85, 5274–5278.

22. Selley, D.E.; Sim, L.J.; Xiao, R.; Liu, Q.; Childers, S.R. μ-Opioid Receptor-Stimulated Guanosine-5′-O-(γ-thio)-triphosphate Binding in Rat Thalamus and Cultured Cell Lines: Signal Transduction Mechanisms Underlying Agonist Efficacy. Molecular Pharmacology 1997, 51, 87–96, doi:10.1080/02701367.2016.1213610.

23. Selley, D.E.; Liu, Q.; Childers, S.R. Signal transduction correlates of mu opioid agonist intrinsic efficacy: receptor-stimulated [35S]GTP gamma S binding in mMOR-CHO cells and rat thalamus. J Pharmacol Exp Ther 1998, 285, 496–505.

24. Lester, P.A.; Traynor, J.R. Comparison of the in vitro efficacy of mu, delta, kappa and ORL1 receptor agonists and non-selective opioid agonists in dog brain membranes. Brain Res 2006, 1073-1074, 290–296, doi:10.1016/j.brainres.2005.12.066.

25. Romero, A.; Miranda, H.F.; Puig, M.M. Analysis of the opioid-opioid combinations according to the nociceptive stimulus in mice. Pharmacol Res 2010, 61, 511–518, doi:10.1016/j.phrs.2010.02.011.

26. Lutfy, K.; Eitan, S.; Bryant, C.D.; Yang, Y.C.; Saliminejad, N.; Walwyn, W.; Kieffer, B.L.; Takeshima, H.; Carroll, F.I.; Maidment, N.T.; et al. Buprenorphine-induced antinociception is mediated by mu-opioid receptors and compromised by concomitant activation of opioid receptor-like receptors. J Neurosci 2003, 23, 10331–10337.

27. Khroyan, T.V.; Wu, J.; Polgar, W.E.; Cami-Kobeci, G.; Fotaki, N.; Husbands, S.M.; Toll, L. BU08073 a buprenorphine analogue with partial agonist activity at mu-receptors in vitro but long-lasting opioid antagonist activity in vivo in mice. Br J Pharmacol 2014, 172, 668–680, doi:10.1111/bph.12796.

28. Acevedo-Canabal, A.; Pantouli, F.; Ravichandran, A.; Rullo, L.; Bohn, L.M. 3.21 - Pharmacological Diversity in Opioid Analgesics: Lessons From Clinically Useful Drugs. In Comprehensive Pharmacology, Kenakin, T., Ed.; Elsevier: Oxford, 2022; pp. 478–493.

29. Lalley, P.M. Opioidergic and dopaminergic modulation of respiration. Respir Physiol Neurobiol 2008, 164, 160–167, doi:10.1016/j.resp.2008.02.004.

30. Torralva, R.; Janowsky, A. Noradrenergic Mechanisms in Fentanyl-Mediated Rapid Death Explain Failure of Naloxone in the Opioid Crisis. J Pharmacol Exp Ther 2019, 371, 453–475, doi:10.1124/jpet.119.258566.

31. Hill, R.; Santhakumar, R.; Dewey, W.; Kelly, E.; Henderson, G. Fentanyl depression of respiration: Comparison with heroin and morphine. Br J Pharmacol 2020, 177, 254–266, doi:10.1111/bph.14860.

32. Kudla, L.; Bugno, R.; Podlewska, S.; Szumiec, L.; Wiktorowska, L.; Bojarski, A.J.; Przewlocki, R. Comparison of an Addictive Potential of μ-Opioid Receptor Agonists with G Protein Bias: Behavioral and Molecular Modeling Studies. Pharmaceutics 2022, 14, doi:https://doi.org/10.3390/pharmaceutics14010055.

