## Supplementary material for "Hyperactivity in mice induced by opioid agonists with partial intrinsic efficacy and biased agonism; alone and in combination with morphine": Sup Figures

### Supplemental Figures

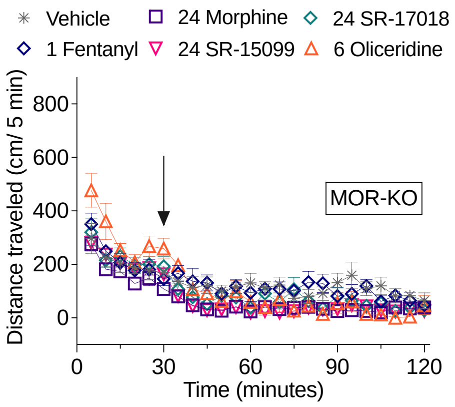


**MOR-KO drug vs vehicle** Two-way RM ANOVA

| Figure | Drug vs vehicle | Time (min) | F (DFn, DFd) Time x Drug | p-Value | F (DFn, DFd) Drug v Vehicle | p-Value | Difference | SE | Drug treatment n |
|---|---|---|---|---|---|---|---|---|---|
| SF1A | 1 Fentanyl | 35-120 | F (17, 204) = 1.284 | P=0.2055 | F (1, 12) = 0.02158 | P=0.8857 | 4.46 | 30.36 | 7 |
| | 24 Morphine | 35-120 | F (17, 272) = 1.151 | P=0.3055 | F (1, 16) = 8.927 | P=0.0087 | -66.85 | 22.37 | 7 |
| | 6 Oliceridine | 35-120 | F (17, 221) = 2.231 | P=0.0044 | F (1, 13) = 2.472 | P=0.1399 | -46.65 | 29.67 | 4 |
| | 24 SR-15099 | 35-60 | F (17, 340) = 1.281 | P=0.2016 | F (1, 20) = 13.60 | P=0.0015 | -68.43 | 18.56 | 11 |
| | 24 SR-17018 | 35-120 | F (17, 221) = 0.8418 | P=0.6436 | F (1, 13) = 1.856 | P=0.1962 | -40.36 | 29.63 | 4 |

vehicle: n=11

**SFigure 1,** ***A lack of opioid-induced hyperactivity in MOR-KO mice.*** Male MOR-KO (n=4-11) were IP treated with vehicle or the highest dose of each opioid tested after a 30-minute habituation period. Distance traveled (cm/5-minute bins) for 90 minutes was recorded and no locomotor stimulation was observed in the MOR-KO. Data are presented as mean ± SEM.

A. Morphine potentiates *d*-Amphetamine LMA

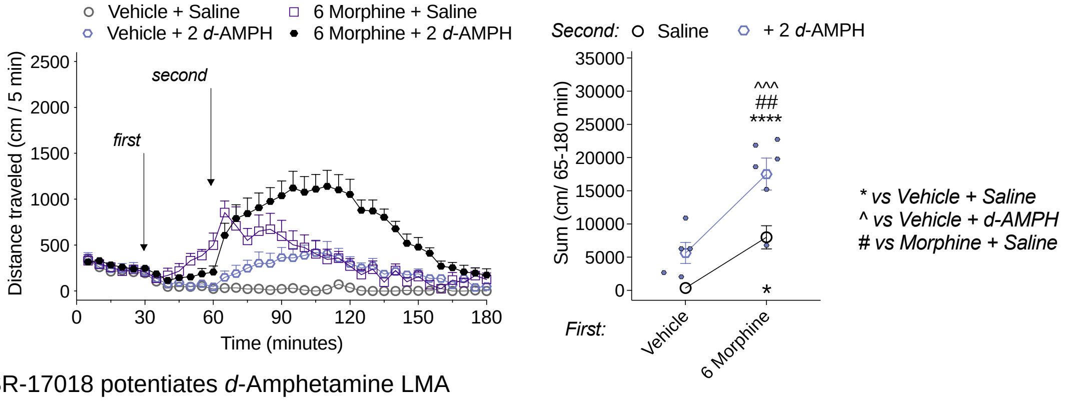


B. SR-17018 potentiates *d*-Amphetamine LMA

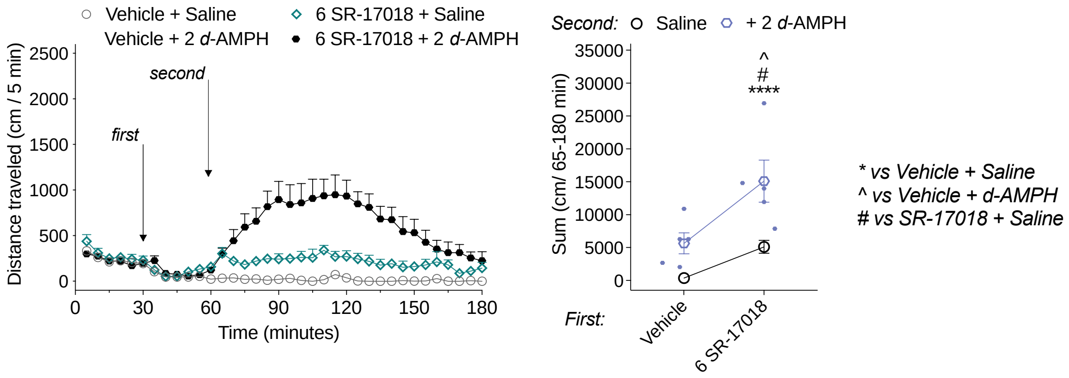


**Locomotor Activity**

**TIMECOURSE** Two-way RM ANOVA

| Figure | Treatment | Compared to | Time | F (DFn, DFd) time x drug | p-Value | F (DFn, DFd) drug effect | p-Value | treatment n |
|---|---|---|---|---|---|---|---|---|
| SF2A | Vehicle + d-Amphetamine | Vehicle + Saline | 65-180 | F (29, 261) = 7.363 | <0.0001 | F (1, 9) = 12.34 | 0.0066 | 5 |
| | 6 Morphine + Saline | Vehicle + Saline | 65-180 | F (29, 232) = 10.67 | <0.0001 | F (1, 8) = 30.45 | 0.0006 | 4 |
| | 6 Morphine + d-Amphetamine | Vehicle + Saline | 65-180 | F (29, 290) = 20.05 | <0.0001 | F (1, 10) = 50.83 | <0.0001 | 5 |
| | Vehicle + d-Amphetamine | 6 Morphine + d-Amphetamine | 65-180 | F (29, 261) = 6.634 | <0.0001 | F (1, 9) = 15.35 | 0.0035 | 5 |
| | 6 Morphine + Saline | 6 Morphine + d-Amphetamine | 65-180 | F (29, 232) = 7.194 | <0.0001 | F (1, 8) = 6.193 | 0.0376 | 5 |
| SF2B | Vehicle + d-Amphetamine | Vehicle + Saline | 65-180 | F (29, 261) = 7.363 | <0.0001 | F (1, 9) = 12.34 | 0.0066 | 8 |
| | 6 SR-17018 + Saline | Vehicle + Saline | 65-180 | F (29, 232) = 4.234 | <0.0001 | F (1, 8) = 44.02 | 0.0002 | 4 |
| | 6 SR-17018 + d-Amphetamine | Vehicle + Saline | 65-180 | F (29, 261) = 14.73 | <0.0001 | F (1, 9) = 27.44 | 0.0005 | 7 |
| | Vehicle + d-Amphetamine | 6 SR-17018 + d-Amphetamine | 65-180 | F (29, 232) = 3.882 | <0.0001 | F (1, 8) = 7.116 | 0.0285 | 4 |
| | 6 SR-17018 + Saline | 6 SR-17018 + d-Amphetamine | 65-180 | F (29, 203) = 5.484 | <0.0001 | F (1, 7) = 7.291 | 0.0306 | 5 |

**SUMS** **Ordinary one-way ANOVA, Šídák's multple comparison post-hoc**

| Figure | Treatment | Compared to | Time | p-Value | treatment n | |
|---|---|---|---|---|---|---|
| SF2A | Vehicle + d-Amphetamine | Vehicle + Saline | 65-180 | 0.1825 | 5 | ns |
| | 6 Morphine + Saline | Vehicle + Saline | 65-180 | 0.0375 | 4 | * |
| | 6 Morphine + d-Amphetamine | Vehicle + Saline | 65-180 | <0.0001 | 5 | **** |
| | Vehicle + d-Amphetamine | 6 Morphine + d-Amphetamine | 65-180 | 0.0005 | 5 | ^^^ |
| | 6 Morphine + Saline | 6 Morphine + d-Amphetamine | 65-180 | 0.0075 | 5 | ## |
| SF2B | Vehicle + d-Amphetamine | Vehicle + Saline | 65-180 | 0.2198 | 8 | ns |
| | 6 SR-17018 + Saline | Vehicle + Saline | 65-180 | 0.3739 | 4 | ns |
| | 6 SR-17018 + d-Amphetamine | Vehicle + Saline | 65-180 | <0.0001 | 7 | **** |
| | Vehicle + d-Amphetamine | 6 SR-17018 + d-Amphetamine | 65-180 | 0.0102 | 4 | ^ |
| | 6 SR-17018 + Saline | 6 SR-17018 + d-Amphetamine | 65-180 | 0.0107 | 5 | # |

**SFigure 2.** ***SR-17018, like, morphine, increases d-amphetamine-induced locomotor activity.*** **A-B**. Pretreatment with the vehicle, morphine (6 mg/kg, i.p.), or SR-17018 (6 mg/kg, i.p.) for 30 minutes followed by challenge with saline or *d*-amphetamine (2 mg/kg, i.p.) in male C57BL6 mice (n=4-6). Effects over time *(left)* and sum of the distance traveled (cm) from 65-180 minutes for individual mice *(right)* are shown in (**A-B**). Comparison of sums are indicated in the figure. Data are presented as mean ± SEM of the total distance (cm).

### A. SR-17018 reversal of Morphine LMA @ 30 min

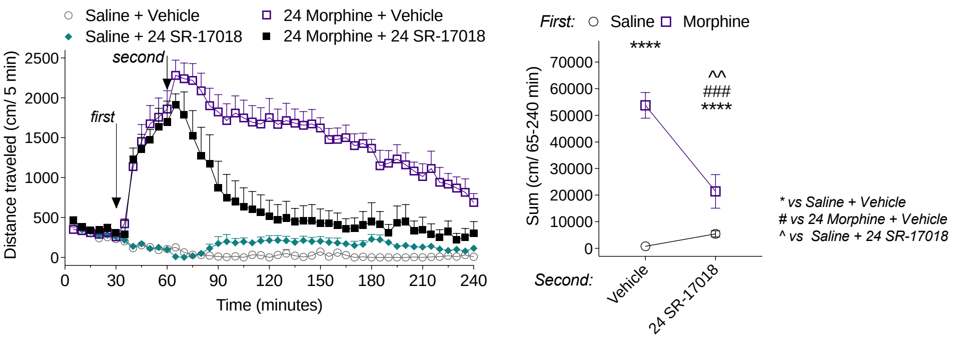


### B. SR-17018 reversal of Morphine LMA @ 60 min

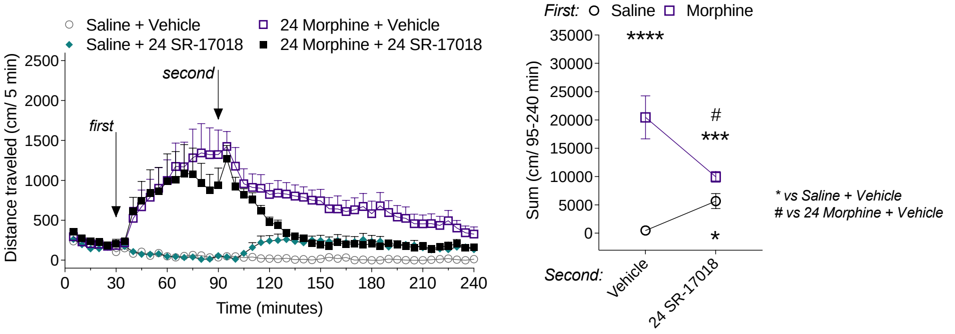


### C. Naloxone reversal of Morphine LMA @ 60 min

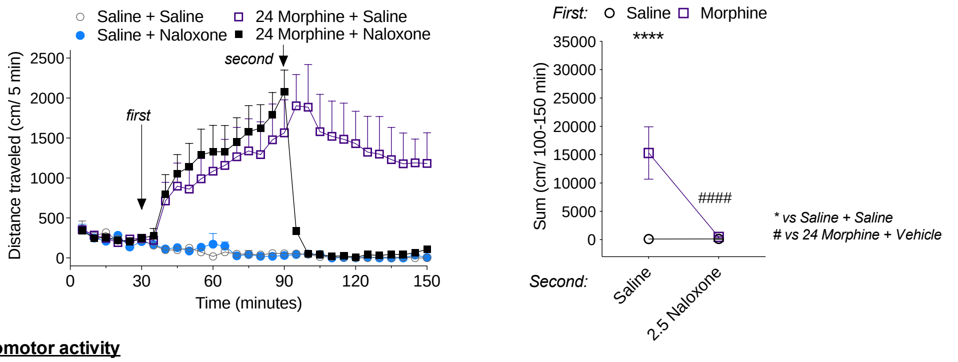


**Locomotor activity**

**Timecourse:** Two-way RM ANOVA

| Figure | Treatment | Compared to | Time | F (DFn, DFd) time x drug | p-Value | F (DFn, DFd) drug effect | p-Value | treatment n |
|---|---|---|---|---|---|---|---|---|
| SF3A | 24 Morphine + Vehicle | Saline + Vehicle | 65-240 | F (35, 350) = 7.374 | <0.0001 | F (1, 10) = 83.36 | <0.0001 | 7 |
| | Saline + 24 SR-17018 | Saline + Vehicle | 65-240 | F (35, 350) = 2.886 | <0.0001 | F (1, 10) = 11.04 | 0.0077 | 7 |
| | Saline + Vehicle | 24 Morphine + 24 SR-17018 | 65-240 | F (35, 280) = 12.94 | <0.0001 | F (1, 8) = 10.67 | 0.0114 | 5 |
| | 24 Morphine + Vehicle | 24 Morphine + 24 SR-17018 | 65-240 | F (35, 350) = 2.192 | 0.0002 | F (1, 10) = 17.14 | 0.002 | 7 |
| | 24 Morphine + 24 SR-17018 | Saline + 24 SR-17018 | 65-240 | F (35, 350) = 20.51 | <0.0001 | F (1, 10) = 8.687 | 0.0146 | 5 |
| SF3B | 24 Morphine + Vehicle | Saline + Vehicle | 95-240 | F (29, 290) = 4.476 | <0.0001 | F (1, 10) = 13.19 | 0.0046 | 7 |
| | Saline + 24 SR-17018 | Saline + Vehicle | 95-240 | F (29, 174) = 3.693 | <0.0001 | F (1, 6) = 15.24 | 0.0079 | 7 |
| | Saline + Vehicle | 24 Morphine + 24 SR-17018 | 95-240 | F (29, 290) = 13.26 | <0.0001 | F (1, 10) = 48.75 | <0.0001 | 5 |
| | 24 Morphine + Vehicle | 24 Morphine + 24 SR-17018 | 95-240 | F (29, 406) = 1.924 | 0.0032 | F (1, 14) = 7.235 | 0.0176 | 7 |
| | 24 Morphine + 24 SR-17018 | Saline + 24 SR-17018 | 95-240 | F (29, 290) = 17.39 | <0.0001 | F (1, 10) = 7.040 | 0.0242 | 5 |
| SF3C | Saline + 2.5 Naloxone | Saline + Saline | 100-150 | F (10, 60) = 0.6125 | 0.7971 | F (1, 6) = 0.3596 | 0.5706 | 4 |
| | 24 morphine + Saline | Saline + Saline | 100-150 | F (10, 80) = 2.870 | 0.0042 | F (1, 8) = 6.920 | 0.0302 | 6 |
| | Saline + Saline | 24 morphine + 2.5 naloxone | 100-150 | F (10, 80) = 1.639 | 0.1106 | F (1, 8) = 6.970 | 0.0297 | 4 |
| | 24 morphine + 2.5 naloxone | 24 morphine + Saline | 100-150 | F (10, 100) = 5.270 | <0.0001 | F (1, 10) = 10.28 | 0.0094 | 6 |
| | 24 morphine + 2.5 naloxone | Saline + 2.5 Naloxone | 100-150 | F (10, 80) = 1.101 | 0.3717 | F (1, 8) = 5.600 | 0.0455 | 6 |

**SUMS Ordinary one-way ANOVA, Šídák's multple comparison post-hoc**

| Figure | Treatment | Compared to | Time | p-Value | treatment n | |
|---|---|---|---|---|---|---|
| SF3A | 24 Morphine + Vehicle | Saline + Vehicle | 65-240 | 0.9313 | 7 | ns |
| | Saline + 24 SR-17018 | Saline + Vehicle | 65-240 | <0.0001 | 7 | **** |
| | Saline + Vehicle | 24 Morphine + 24 SR-17018 | 65-240 | 0.014 | 5 | * |
| | 24 Morphine + Vehicle | 24 Morphine + 24 SR-17018 | 65-240 | 0.0491 | 7 | * |
| | 24 Morphine + 24 SR-17018 | Saline + 24 SR-17018 | 65-240 | <0.0001 | 5 | **** |
| SF3B | 24 Morphine + Vehicle | Saline + Vehicle | 95-240 | 0.0004 | 7 | *** |
| | Saline + 24 SR-17018 | Saline + Vehicle | 95-240 | 0.8083 | 7 | ns |
| | Saline + Vehicle | 24 Morphine + 24 SR-17018 | 95-240 | 0.1414 | 5 | ns |
| | 24 Morphine + Vehicle | 24 Morphine + 24 SR-17018 | 95-240 | 0.0235 | 7 | * |
| | 24 Morphine + 24 SR-17018 | Saline + 24 SR-17018 | 95-240 | 0.8354 | 5 | ns |
| SF3C | Saline + 2.5 Naloxone | Saline + Saline | 100-150 | >0.9999 | 4 | ns |
| | 24 morphine + Saline | Saline + Saline | 100-150 | 0.0093 | 6 | ** |
| | Saline + Saline | 24 morphine + 2.5 naloxone | 100-150 | >0.9999 | 4 | ns |
| | 24 morphine + 2.5 naloxone | 24 morphine + Saline | 100-150 | 0.0046 | 6 | ** |
| | 24 morphine + 2.5 naloxone | Saline + 2.5 Naloxone | 100-150 | >0.9999 | 6 | ns |

**SFigure 3. *SR-17018 reversal of morphine-induced locomotor activity*.** The basal activity was recorded for 30 minutes followed by 30- (**A**) or 60-minute (**B**) treatment with saline or morphine (24 mg/kg, i.p.) before vehicle or SR-17018 (24 mg/kg, i.p.) challenge in male C57BL6 mice (n=4-8). Effects over time *(left)* and sum or the distance traveled (cm) from 65-240 minutes for individual mice *(right)* are shown in (**A-B**). **C**. Naloxone (2.5 mg/kg, i.p.) reversal after 60 minutes of morphine locomotor activity is shown for reference (n = 4-6). Comparison of sums are indicated in the figure. Sums figures have different Y-axis scales based on the maximal stimulation per treatment combination and we are showing individual sums per mouse. **D**. Table showing least square mean differences Tukey HSD analysis for pretreatment x second treatment interactions after saline or morphine + vehicle, SR-17018, or naloxone challenge. Data are presented as mean ± SEM of the total distance (cm).

A. Locomotor Activity: SR-15099 + Morphine

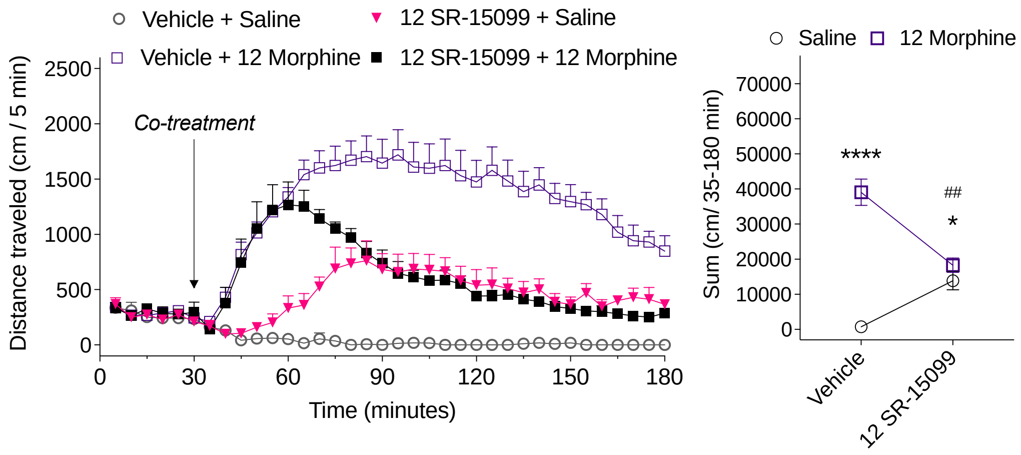


**Locomotor activity**

**Timecourse:** Two-way RM ANOVA

| Figure | Treatment | Compared to | Time | F (DFn, DFd) (Minutes x Days) | p-Value | F (DFn, DFd) (Day 1 v Day 7) | p-Value | treatment n |
|---|---|---|---|---|---|---|---|---|
| SF5 | Vehicle + 12 Morphine | Vehicle + Saline | 35-180 | F (29, 319) = 5.916 | <0.0001 | F (1, 11) = 44.41 | <0.0001 | 4 |
| | 12 SR-15099 + Saline | Vehicle + Saline | 35-180 | F (29, 174) = 8.913 | <0.0001 | F (1, 6) = 25.76 | 0.0023 | 4 |
| | 12 SR-15099 + 12 Morphine | Vehicle + Saline | 35-180 | F (29, 174) = 15.95 | <0.0001 | F (1, 6) = 74.92 | 0.0001 | 4 |
| | Vehicle + 12 Morphine | 12 SR-15099 + 12 Morphine | 35-180 | F (29, 319) = 4.950 | <0.0001 | F (1, 11) = 12.43 | 0.0047 | 9 |
| | 12 SR-15099 + 12 Morphine | 12 SR-15099 + Saline | 35-180 | F (29, 174) = 11.98 | <0.0001 | F (1, 6) = 1.830 | 0.2249 | 4 |

**SUMS Ordinary one-way ANOVA, Šídák's multple comparison post-hoc**

| Figure | Treatment | Compared to | Time | p-Value | treatment n | |
|---|---|---|---|---|---|---|
| SF5 | Vehicle + 12 Morphine | Vehicle + Saline | 35-180 | <0.0001 | 4 | **** |
| | 12 SR-15099 + Saline | Vehicle + Saline | 35-180 | 0.1695 | 4 | ns |
| | 12 SR-15099 + 12 Morphine | Vehicle + Saline | 35-180 | 0.0366 | 4 | * |
| | Vehicle + 12 Morphine | 12 SR-15099 + 12 Morphine | 35-180 | 0.0028 | 9 | ** |
| | 12 SR-15099 + 12 Morphine | 12 SR-15099 + Saline | 35-180 | 0.9513 | 4 | ns |

**SFigure 4** ***Co-treatment of morphine with SR-15099 attenuates morphine-induced hyperactivity.*** *In vivo* drug interactions in the open field locomotor activity assay showing total distance traveled (cm/ 5-minutes) (*left)* with total distance sums after co-treatment (cm/ 35-180 minutes) (*right*) in male C57BL/6 (n = 4-7). Locomotor activity after cotreatment with vehicle or SR-15099 (12 mg/kg, i.p.) and saline or morphine (12 mg/kg, i.p.). SR-15099 and morphine drug combination attenuate morphine's hyperactivity to the level of SR-15099. Comparison of sums are shown in the figure. Data are presented as mean ± SEM.

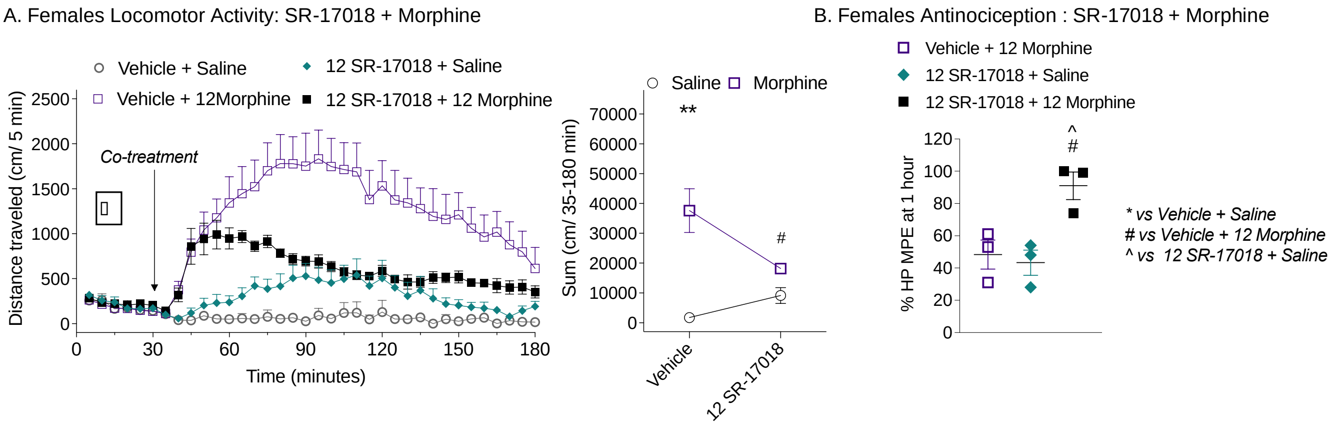


**Locomotor Activity**

**TIMECOURSE** **Two-way RM ANOVA**

| Figure | Treatment | Compared to | Time | F (DFn, DFd) time x drug | p-Value | F (DFn, DFd) drug effect | p-Value | treatment n |
|---|---|---|---|---|---|---|---|---|
| SF4a | Vehicle + 12 Morphine | Vehicle + Saline | 35-180 | F (29, 203) = 6.199 | P<0.0001 | F (1, 7) = 11.12 | P=0.0125 | 6 |
| | 12 SR-17018 + Saline | Vehicle + Saline | 35-180 | F (29, 145) = 2.521 | P=0.0002 | F (1, 5) = 5.285 | P=0.0699 | 4 |
| | 12 SR-17018 + 12 Morphine | Vehicle + Saline | 35-180 | F (29, 290) = 7.029 | P<0.0001 | F (1, 10) = 6.869 | P=0.0256 | 6 |
| | Vehicle + 12 Morphine | 12 SR-17018 + 12 Morphine | 35-180 | F (29, 290) = 7.029 | P<0.0001 | F (1, 10) = 6.869 | P=0.0256 | 6 |
| | 12 SR-17018 + 12 Morphine | 12 SR-17018 + Saline | 35-180 | F (29, 232) = 4.238 | P<0.0001 | F (1, 8) = 13.14 | P=0.0067 | 6 |

**Sums** **Ordinary one-way ANOVA, Šídák's multple comparison post-hoc**

| Figure | Treatment | Compared to | Time | p-Value | treatment n | |
|---|---|---|---|---|---|---|
| SF4a | Vehicle + 12 Morphine | Vehicle + Saline | 35-180 | 0.0014 | 6 | ** |
| | 12 SR-17018 + Saline | Vehicle + Saline | 35-180 | 0.9099 | 4 | ns |
| | 12 SR-17018 + 12 Morphine | Vehicle + Saline | 35-180 | 0.216 | 6 | ns |
| | Vehicle + 12 Morphine | 12 SR-17018 + 12 Morphine | 35-180 | 0.0342 | 6 | * |
| | 12 SR-17018 + 12 Morphine | 12 SR-17018 + Saline | 35-180 | 0.699 | 6 | ns |

vehicle+saline n=3

**Hot Plate Latency**

**Ordinary one-way ANOVA, Šídák's multple comparison post-hoc**

| | Treatment | Compared to | Time | p-Value | treatment n | |
|---|---|---|---|---|---|---|
| SF4b | Vehicle + 12 Morphine | 12 SR-17018 + Saline | 1h | 0.90960 | 3 | |
| | 12 SR-17018 + 12 Morphine | Vehicle + 12 Morphine | 1h | 0.0274 | 3 | # |
| | 12 SR-17018 + 12 Morphine | 12 SR-17018 + Saline | 1h | 0.017 | 3 | ^ |

**SFigure 5. *In female mice, co-treatment of morphine with SR-17018 decreases morphine-induced hyperactivity without attenuating antinociception.*** Results recapitulate in female C57BL6 mice testing for SR-17018 + morphine locomotor (n = 3-6) and antinociception (n=3). Doses are indicated as mg/kg, i.p. in the figure legends. Data are presented as mean ± SEM.
